# Expansion of the MutS Gene Family in Plants

**DOI:** 10.1101/2024.07.17.603841

**Authors:** Daniel B. Sloan, Amanda K. Broz, Shady A. Kuster, Viraj Muthye, Alejandro Peñafiel-Ayala, Jennifer R. Marron, Dennis V. Lavrov, Luis G. Brieba

## Abstract

The *MutS* gene family is distributed across the tree of life and is involved in recombination, DNA repair, and protein translation. Multiple evolutionary processes have expanded the set of *MutS* genes in plants relative to other eukaryotes. Here, we investigate the origins and functions of these plant-specific genes. Land plants, green algae, red algae, and glaucophytes share cyanobacterial-like *MutS1* and *MutS2* genes that presumably were gained via plastid endosymbiotic gene transfer. *MutS1* was subsequently lost in some taxa, including seed plants, whereas *MutS2* was duplicated in Viridiplantae (i.e., land plants and green algae) with widespread retention of both resulting paralogs. Viridiplantae also have two anciently duplicated copies of the eukaryotic *MSH6* gene (i.e., *MSH6* and *MSH7*) and acquired *MSH1* via horizontal gene transfer – potentially from a nucleocytovirus. Despite sharing the same name, “plant *MSH1*” is not directly related to the gene known as *MSH1* in some fungi and animals, which may be an ancestral eukaryotic gene acquired via mitochondrial endosymbiosis and subsequently lost in most eukaryotic lineages. There has been substantial progress in understanding the functions of *MSH1* and *MSH6*/*MSH7* in plants, but the roles of the cyanobacterial-like *MutS1* and *MutS2* genes remain uncharacterized. Known functions of bacterial homologs and predicted protein structures, including fusions to diverse nuclease domains, provide hypotheses about potential molecular mechanisms. Because most plant-specific MutS proteins are targeted to the mitochondria and/or plastids, the expansion of this family appears to have played a large role in shaping plant organelle genetics.

**One-Sentence Summary:** Plants are distinguished from other eukaryotes by a functionally diverse complement of MutS proteins gained via a combination of gene duplication, endosymbiotic gene transfer, and horizontal gene transfer.

## Introduction

*MutS* genes comprise an ancient family with representatives in bacteria, archaea, eukaryotes, and even viruses (Figure 1) (Ogata et al. 2011). Phylogenetic approaches have been central to dissecting the history of gene duplication, horizontal transfer, and diversification that has defined the composition of the *MutS* family (Culligan et al. 2000; Lin et al. 2007; Bilewitch and Degnan 2011; Hofstatter and Lahr 2021). Indeed, the term “phylogenomics” was coined in the context of analyzing *MutS* evolutionary history (Eisen et al. 1997; Eisen 1998).

**Figure 1.**
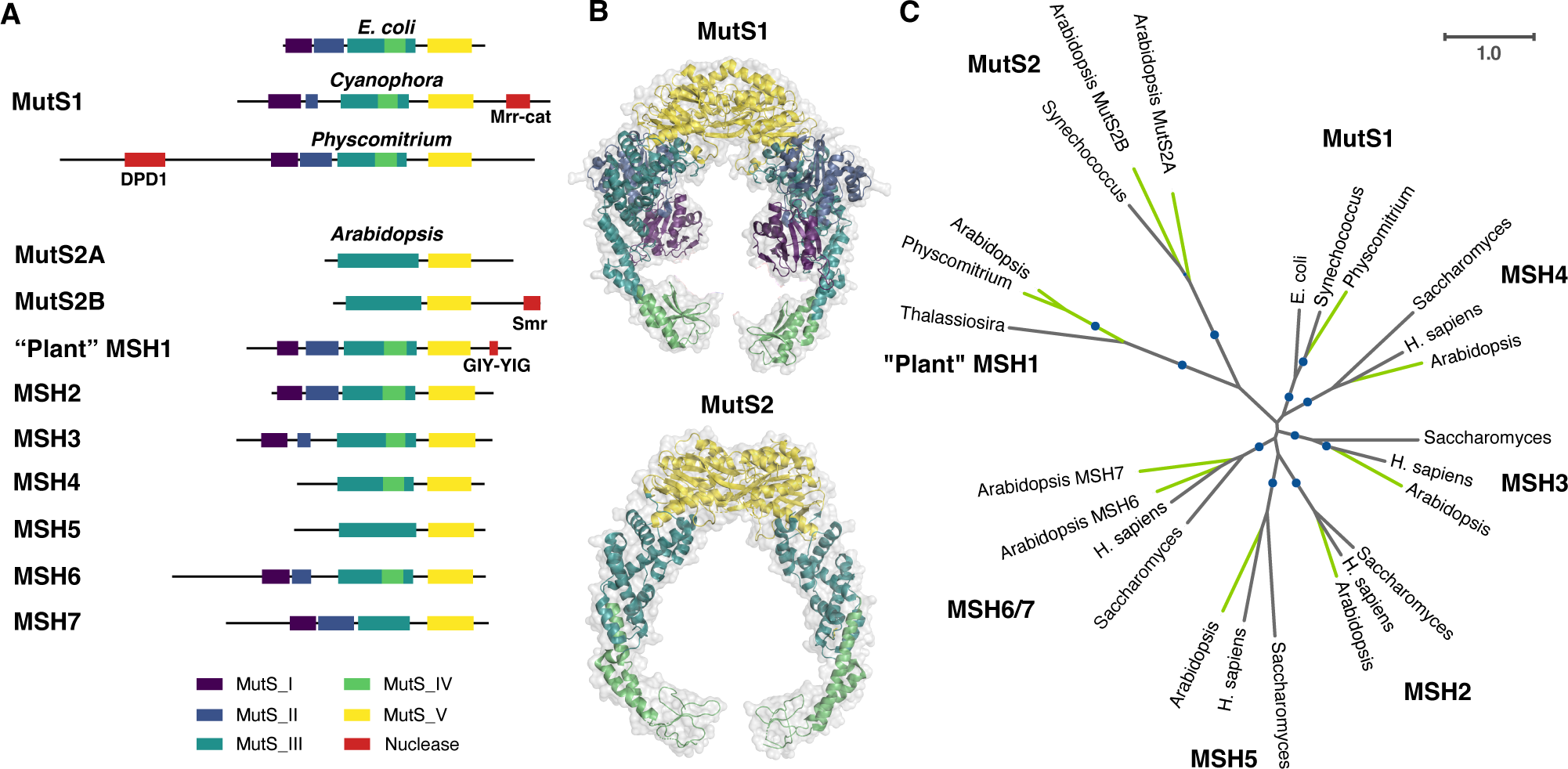
The plant MutS protein family. (A) Summary of domain architecture for representatives of each of the eight MutS subfamilies that are found in plants, highlighting fusions with different predicted nuclease domains. *Arabidopsis* proteins are shown as representatives for each MutS type except MutS1 for which the moss *Physcomitrium* is used because *Arabidopsis* and other seed plants appear to have lost the cyanobacterial-like MutS1 that was likely acquired from plastids. Domain annotation was based on InterProScan except that Domains II-IV were not detected in *Arabidopsis* MSH1 and were added manually based on sequence alignment and structural homology and that the Mrr_cat domain was predicted by CD-Search Tool. Despite the identified similarity to an Mrr_cat domain, actual nuclease activity may be unlikely (see main text). (B) Structures of bacterial MutS1 (*E. coli* PDB: 7AI5) and bacterial MutS2 (*Thermus thermophilus* PDB: 7VUF) dimers, showing examples of MutS enzymes both with (MutS1) and without (MutS2) the N-terminal mismatch recognition and connector domains. Color coding reflects domain architecture shown in Panel A. Bacterial MutS1 is divided into five defined domains: mismatch binding domain (Domain I, residues 1–115, deep purple), connector domain (Domain II, residues 116−266, blue), core domain (Domain III, 267−443 and 504−567, teal), clamp and levers domain (Domain IV, 444−503, green) and ATPase domain plus helix-turn-helix domain (Domain V, 568−765 and 765−800, yellow) (Bhairosing-Kok 2021; Fernandez-Leiro et al. 2021). Bacterial MutS2 is divided into three defined domains: core domain (Domain III, residues 1−131 and 248−277, teal), clamp and levers domain (Domain IV, residues 132−247, green), and ATPase domain plus helix-turn-helix domain (Domain V, residues 248 to 486, yellow) (Fukui et al. 2022). The absence of domain I and II in bacterial MutS2 creates a DNA binding site of more than 70 Å able to accommodate Holliday junctions or D-loops (Fukui et al. 2022). (C) Unrooted maximum-likelihood phylogenetic tree showing each of the eight MutS subfamilies in which plant representatives have been identified. Plant lineages are highlighted in green. Clades with bootstrap support >80% are indicated with circles. See Supplementary Text S1 for information on tree reconstruction, domain prediction, and visualization of protein structures.

The function of *MutS* genes was initially studied with respect to bacterial *MutS1* (often simply referred to as *MutS*) and the DNA mismatch repair (MMR) pathway (Siegel and Bryson 1963; Rydberg 1978; Su and Modrich 1986). The general role of MutS proteins in this pathway is to scan DNA and recognize structural distortions resulting from bases that are damaged, improperly paired (mismatches from polymerase errors), or unpaired (indels from polymerase slippage), at which point additional proteins (e.g., MutL) are recruited to initiate DNA repair (Li 2008). However, many MutS family members lack MMR function and instead play roles in recombination and meiosis (Eisen 1998). It was also recently discovered that members of the bacterial MutS2 subfamily rescue stalled and collided ribosomes during protein translation, meaning that some MutS proteins have functions that are entirely unrelated to DNA interactions (Cerullo et al. 2022; Park et al. 2024). Furthermore, some of the major MutS subfamilies have yet to be characterized at all (Ogata et al. 2011), so it is likely that MutS proteins are involved in an even more diverse array of functions.

MutS proteins also vary substantially in domain architecture (Figure 1) (Ogata et al. 2011). The ATPase domain (Domain V, pfam00488) is found in all subfamilies and maintains the highest levels of amino-acid sequence conservation across the family (Sachadyn 2010). The DNA-binding clamp and levers (Domain IV, pfam05190) and the core (Domain III, pfam05192) are also present in most MutS subfamilies. In contrast, the N-terminal mismatch recognition domain (Domain I, pfam01624) and adjacent connector (Domain II, pfam05188) are missing from most of the MutS subfamilies that do not function in MMR (Lamers et al. 2000). For instance, MutS2 lacks Domains I and II, resulting in a wider DNA binding site that allows interactions with voluminous DNA structures like Holliday junctions or D-loops (Lin et al. 2007; Fukui et al. 2008, 2022). In addition, MutS protein diversity reflects the frequent acquisition of novel domains, including many with predicted nuclease function (Ogata et al. 2011). Despite these variations in domain architecture, MutS proteins share a tendency to assemble as dimers or tetramers (Mendillo et al. 2007). Bacterial MutS proteins (e.g., MutS1) typically assemble with themselves to form homodimers (Lamers et al. 2000). In contrast, eukaryotes have many MutS Homolog (MSH) proteins (Kolodner 1996), which can pair in various combinations as heterodimers (Snowden et al. 2004; Li 2008).

Here, our goal is to characterize the expansionary history of the MutS gene family in plants relative to other eukaryotes. We investigate the origins of plant-specific MutS genes, exploring functions that have already been determined for some of these genes and pointing to key knowledge gaps and testable hypotheses for family members that are currently uncharacterized.

## The Typical Eukaryotic Set of *MutS* Genes

Early studies in yeast (*Saccharomyces cerevisiae*) identified a set of six *MutS* homologs that were termed *MSH1-MSH6* (Kolodner 1996). Five of these yeast genes (*MSH2-MSH6*) have orthologs in mammals and plants (Eisen 1998; Culligan and Hays 2000; Lin et al. 2007). Phylogenetic analysis has indicated that these five genes represent an ancestral eukaryotic set resulting from sequential duplications of a gene that was inherited from the Asgard archaeal progenitor of the eukaryotic host cell (Hofstatter and Lahr 2021). However, there is limited statistical support for this inferred archaeal relationship. A lack of robust phylogenetic resolution is an unfortunate recurring theme in inferring the history of the *MutS* family. Because many of the diversification events in this family span deep splits in the tree of life, it is challenging to confidently reconstruct these relationships from a single protein alignment, and alternative evolutionary scenarios have been proposed (Eisen 1998; Culligan et al. 2000; Lin et al. 2007; Rossier et al. 2024). In contrast to the other five yeast *MSH* genes, *MSH1* lacks orthologs in many eukaryotic lineages, including most animals and plants. Instead, the yeast *MSH1* gene is similar to the bacterial *MutS1* gene (Ogata et al. 2011), and eukaryotic orthologs have only been previously identified in other fungi, some non-bilaterian animals, and the amoebozoan *Dictyostelium* (Lin et al. 2007; Muthye and Lavrov 2021). *MSH1* is also the only of the six yeast *MSH* genes that encodes a mitochondrially targeted protein, as the other five all have roles in the nucleus.

In contrast to bacterial MutS proteins, which form homodimers, the nuclear-localized MSH proteins in eukaryotes function as obligate heterodimer complexes, which dictates their roles in MMR or in processing recombination intermediates. MSH2 can pair with MSH6 to form a heterodimer known as MutSα or with MSH3 to form the MutSβ heterodimer. These complexes function in MMR, although they have different DNA lesion targets (e.g., mismatches vs. indel loops) (Culligan and Hays 2000; Li 2008). In contrast, MSH4 and MSH5 heterodimerize with each other and mediate homologous crossovers in meiosis (Ross-Macdonald and Roeder 1994; Higgins et al. 2004; Snowden et al. 2004). As the sole yeast MutS protein localized to the mitochondria, MSH1 presumably assembles as a homodimer, and it has been found to play a key role in mitochondrial DNA repair and maintenance (Chi and Kolodner 1994; Mookerjee and Sia 2006; Pogorzala et al. 2009).

The similarity in the *MutS* gene set between yeast and animals shows that there can be a high degree of stability and conservation in this gene family, even between deeply divergent lineages. However, the evolution of the *MutS* family has been dynamic in the broader “plant” (Archaeplastida) lineage, including an expansion in the size of the family through a combination of gene duplication, endosymbiotic gene transfer (EGT), and horizontal gene transfer (HGT). Here, we discuss the origins and (potential) functions of each of the plant-specific *MutS* genes.

## Viridiplantae Contain Ancient *MSH6/MSH7* Duplicates with Divergent Functions in Nuclear Genome Maintenance

Initial searches for homologs of the yeast and mammal *MSH2*-*MSH6* gene set revealed that plants contained two *MSH6*-like genes (AT4G02070 and AT3G24495 in *Arabidopsis*). Accordingly, this gene pair was referred to as either *MSH6-1/MSH6-2* (Adé et al. 1999) or *MSH6/MSH7* (Culligan and Hays 2000), with the latter having since been adopted as the more common nomenclature (Lin et al. 2007; Tam et al. 2009). Despite both being *MSH6* homologs, these genes are substantially divergent in sequence from each other (less than 40% amino acid identity between the two *Arabidopsis* copies), implying an ancient duplication followed by long-term retention of both paralogs.

A still-unresolved question that arose from the discovery of plant *MSH6* and *MSH7* is whether these duplicates specifically emerged in the plant lineage or are products of an even earlier duplication with subsequent gene losses in other eukaryotic lineages. Phylogenetic analysis indicates that the duplication must have occurred before the most recent common ancestor (MRCA) of Viridiplantae because *MSH6* and *MSH7* form well-supported monophyletic clades with representatives spanning the diversity of this taxonomic group (Figure 2a). It is also possible that the duplication event was even more ancient. Indeed, the gene tree topology supports a complex history of multiple ancient duplications and differential gene losses because of conflicts with known species relationships (e.g., animal *MSH6* identified as sister to plant *MSH6* rather than in a clade with fungal *MSH6*). Despite substantial statistical (bootstrap) support for some of these relationships, we suggest caution in inferring that the duplication(s) occurred prior to the divergence of major eukaryotic supergroups. Variation in evolutionary rates can bias inferred tree topologies due to long-branch attraction, and the plant *MSH7* gene appears to have evolved at a higher rate than its *MSH6* counterpart (Figure 2a). The absence of duplicates in the other sampled eukaryotic lineages (animals, fungi, and red algae) could be explained by the inferred tree topology being erroneous and a duplication having occurred after the Viridiplantae lineage split off from the rest of the tree of life. Under either scenario, however, extant plants are distinct in that they have two different *MSH6*-like genes.

**Figure 2.**
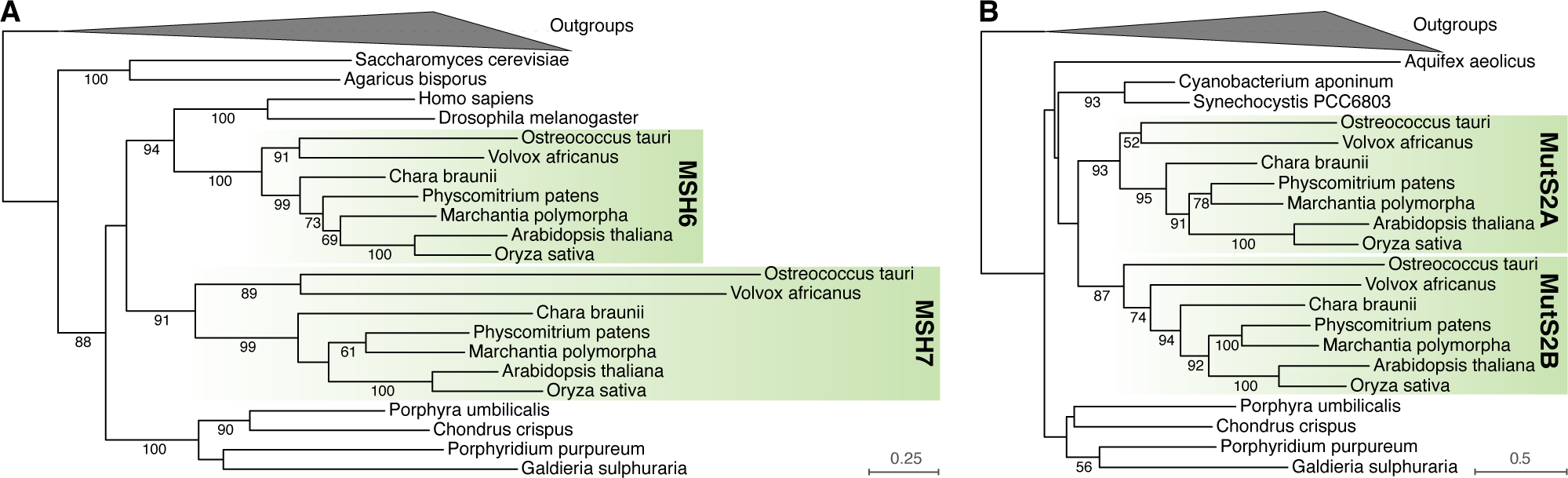
Viridiplantae maintain ancient duplicates of the *MSH6* and *MutS2* genes. Maximum-likelihood phylogenetic trees based on sequences of (A) MSH6 and (B) MutS2 proteins. In both cases, two ancient gene copies are conserved across all sampled green algae and land plants (highlighted clades). Bootstrap values are reported for clades with >50% support. MutS1 and MSH3 sequences were used as outgroups to root the MSH6 tree, and MutS1 and MSH5 were used as outgroups to root the MutS2 tree. See Supplementary Text S1 for information on tree reconstruction.

*Arabidopsis* MSH6 and MSH7 are thought to function in the nucleus and have both been found to heterodimerize with MSH2, which is consistent with the dimerization activity of MSH6 in other eukaryotes (Culligan and Hays 2000). Analysis of DNA binding specificities suggests that the MSH2⋅MSH6 heterodimer (MutSα) has preferences for G/T base-pair mismatches and short indels that are similar to those of MutSα in other eukaryotes, whereas the MSH2⋅MSH7 heterodimer may specialize in the recognition of other mismatches and DNA lesions (Wu et al. 2003). The higher rate of accumulation of cyclobutane pyrimidine dimers in response to UV light in *Arabidopsis msh7* mutants relative to wild-type suggests a role in recognition and repair of UV damage (Lario et al. 2015). In addition, MSH7 appears to regulate and suppress recombination, as angiosperm *msh7* mutants exhibit altered rates of homologous and/or homoeologous crossovers in meiosis (Lario et al. 2015; Serra et al. 2021). *Arabidopsis msh7* mutants exhibit accelerated germination, root growth, and production of cauline leaves and axillary inflorescences, which has been attributed to disruption in cell-cycle checkpoints due to inadequate DNA repair (Chirinos-Arias and Spampinato 2020).

A distinctive feature of plant MSH6 relative to its counterparts from other eukaryotes is the presence of a histone reader Tudor domain that binds to H3K4me1 histone modifications and preferentially recruits MMR machinery to gene bodies in the plant nuclear genome (Quiroz et al. 2024). As a result, genic regions experience lower mutation rates than the rest of the genome (Belfield et al. 2018; Monroe et al. 2022). The function of this Tudor domain appears to be analogous to the PWWS domain that recruits mammalian MSH6 to gene bodies via H3K36me3 histone modifications (Quiroz et al. 2024). In contrast, plant MSH7 lacks the Tudor domain and has a highly reduced clamp domain, which has been hypothesized to alter its DNA binding specificity and may be related to the role of MSH7 in regulating recombination (Wu et al. 2003). Thus, although many of the details of plant MSH6 and MSH7 function remain to be elucidated, these ancient duplicates have clearly diverged in protein structure and function to play differentiated roles in nuclear genome maintenance.

“Plant” *MSH1* Functions in Mitochondrial and Plastid Genome Maintenance and was Acquired by Viridiplantae via Horizontal Transfer

*MSH1* is one of the most intriguing and long-studied genes involved in plant mitochondrial and plastid genetics. This nuclear gene was originally identified based on a mutation in *Arabidopsis* that elicited leaf variegation phenotypes that subsequently exhibited patterns of maternal inheritance (Redei 1973). Accordingly, the nuclear gene was named *CHM* for “chloroplast mutator” under the expectation that it was necessary for preventing mutations in the plastid genome. However, subsequent work showed that some of the observed phenotypic effects in *chm* mutants were mediated by structural instability in the mitochondrial genome (Martinez-Zapater et al. 1992; Sakamoto et al. 1996; Abdelnoor et al. 2003; Davila et al. 2011; Zou et al. 2022). When the *Arabidopsis CHM* locus (AT3G24320) was cloned, it was found to encode a MutS homolog that is targeted to mitochondria (later discovered to be dual-targeted to plastids as well), and the gene was renamed *MSH1* as a presumed ortholog of the mitochondrial-targeted *MSH1* gene in yeast (Abdelnoor et al. 2003). However, phylogenetic analysis has since shown that these genes are derived from entirely different parts of the MutS family (Lin et al. 2007; Ogata et al. 2011).

As shown in Figure 3a, the yeast *MSH1* gene forms a clade with bacterial *mutS1* genes and homologs from eukaryotes, including other fungi, some non-bilaterian animals (hexacorals, sponges, and placozoans), amoebozoans (dictyosteliids), stramenopiles (oomycetes and *Hondaea*), and discobids (*Andalucia* and *Naegleria*) (Lin et al. 2007; Bilewitch and Degnan 2011; Ogata et al. 2011; Muthye and Lavrov 2021). Multiple observations support the inference that these genes are mitochondrial in origin and, thus, derived from a gene that was present in the MRCA of all eukaryotes: 1) the eukaryotic genes in this group appear to be most similar to homologs in Alphaproteobacteria – the group from which mitochondria are thought to have derived (Lin et al. 2007; Hofstatter and Lahr 2021); 2) the proteins they encode are often known or predicted to localize to mitochondria (Table 1); 3) they are present in at least three different eukaryotic supergroups (Amorphea, TSAR, and Discoba; Burki et al. 2020). This interpretation implies a history of numerous independent losses given the apparent absence of this gene in most eukaryotic lineages. Such a history seems plausible because the extensive sampling within animals has shown there have been multiple losses even within this one lineage (Muthye and Lavrov 2021). However, alternative scenarios involving more recent HGT(s) from bacteria to eukaryotes and/or subsequent HGT among eukaryotes are also important to consider. For example, the presence of an alphaproteobacterial-like MutS1 in a single genus of red algae (*Galdieria*) is a candidate for an independent HGT event given its absence from other Archaeplastida and the history of extensive HGT in this genus (Qiu et al. 2013; Pandey et al. 2017; Rossoni et al. 2019).

**Figure 3.**
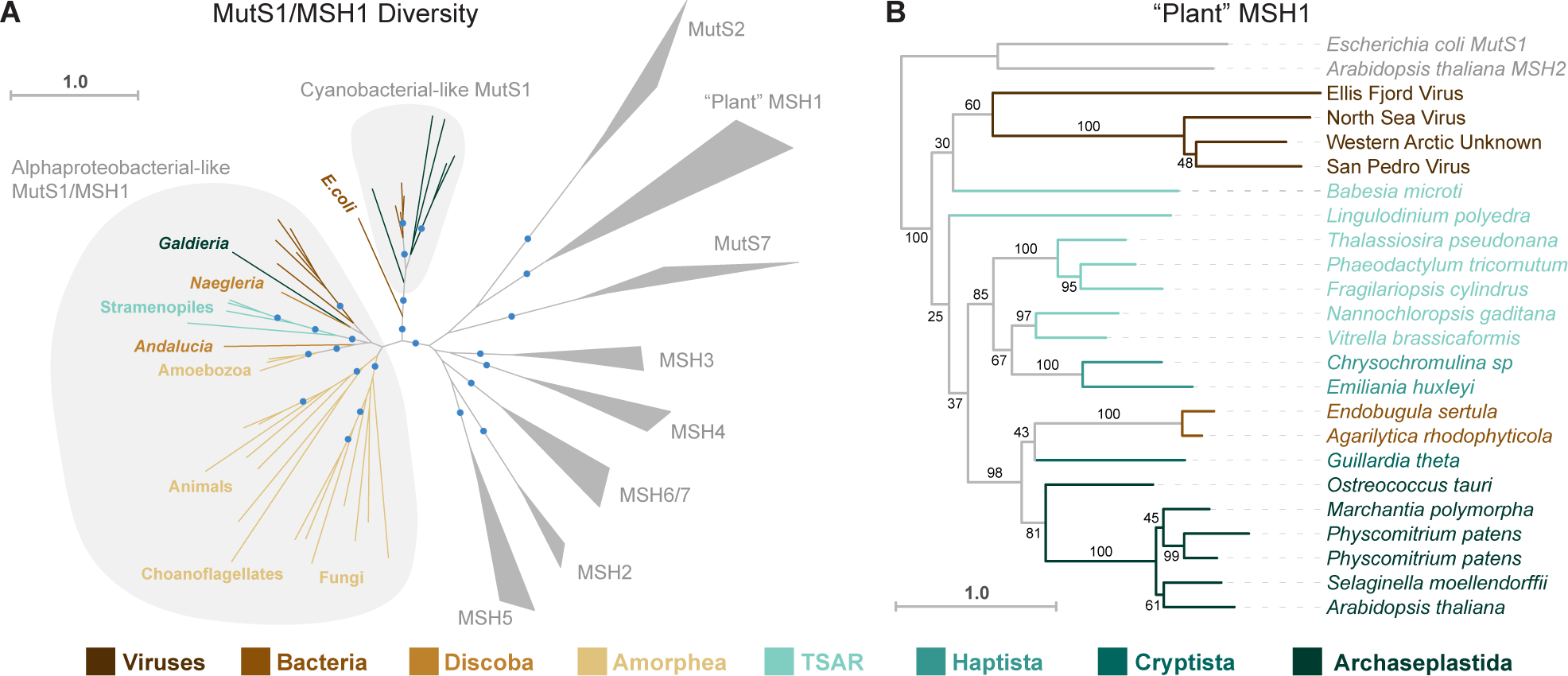
Origins and diversity of MutS proteins with mitochondrial and/or plastid function in eukaryotes. (A) Maximum-likelihood phylogeny with representatives of the MutS subfamilies found in eukaryotes. Sampling emphasizes bacterial-like MutS1 proteins that include the mitochondrial-targeted MSH1 protein that has been described in yeast and other eukaryotes. Representatives of other MutS subfamilies are collapsed into clades (triangles). The indicated grouping of alphaproteobacterial-like MutS1/MSH1 proteins includes representative from multiple eukaryotic supergroups: Amorphea (Amoebozoa, animals, choanoflagellates, and fungi), TSAR (stramenopiles), Discoba (*Andalucia* and *Naegleria*), and Archaeplastida (with the genus *Galdieria* being the only identified representative). The clade of bacteria within this encircled group are all Alphaproteobacteria. This analysis did not recover a single MSH1 clade that was nested within the Alphaproteobacteria or that was monophyletic and sister to the Alphaproteobacteria, but that may reflect distortions from long branches and other artefacts. The eukaryotes within this group generally exhibit strongest similarity to Alphaproteobacteria when searched against all bacteria in the NCBI nr database with BLASTP, and previous phylogenetic analyses with more extensive sampling of bacterial diversity have also recovered affinities for Alphaproteobacteria (Hofstatter and Lahr 2021). An independent plastid-derived origin of MutS1-like proteins in eukaryotes is also indicated, as the encircled group shows representatives from Cyanobacteria and Archaeplastida (see Figure 4 for details). The tree also shows that these MutS1-like proteins are not directly related to the organelle-targeted “plant” MSH1 protein or to MutS7 proteins, which are found in mitochondria of octocorals, as well as some nucleocytoviruses and Epsilonproteobacteria. Clades with bootstrap support >80% are indicated with circles. (B) Maximum-likelihood phylogeny of the “plant” MSH1 clade shows that these proteins are found in lineages outside of Viridiplantae, including various protists, a small clade of Gammaproteobacteria, and some nucleocytoviruses. This disjunct phylogenetic distribution is clear evidence of HGT, but reconstruction of the specific ancient transfer events is challenging given the level of resolution for deep nodes in the tree (numerical values indicate bootstrap support) and the potential for long-branch artefacts. MutS1 and MSH2 representatives were used to root the tree. See Supplementary Text S1 for information on tree reconstruction, domain prediction, and visualization of protein structures.

**Table 1.** Probability of organellar targeting as predicted by TargetP for eukaryotic MSH1 proteins related to alphaproteobacterial MutS1. Reported values are for mitochondrial targeting based on predictions run in “Non-Plant” mode with the exception of *Galdieria*, which was run in “Plant” mode and reports the sum of predicted mitochondrial and plastid targeting.

| Species | Accession | Supergroup/Lineage | Targeting |
| --- | --- | --- | --- |
| *Acytostelium subglobosum* | XP_012750667.1 | Amorphea/Amoebozoa | 0.97 |
| *Cavenderia fasciculata* | XP_004351969.1 | Amorphea/Amoebozoa | 0.00 |
| *Dictyostelium discoideum* | Q552L1 | Amorphea/Amoebozoa | 0.00 |
| *Actinia tenebrosa* | A0A6P8IL16 | Amorphea/Animals | 0.79 |
| *Amphimedon queenslandica* | A0A1X7U1M9 | Amorphea/Animals | 0.65 |
| *Pocillopora damicornis* | A0A3M6TFP8 | Amorphea/Animals | 0.28 |
| *Trichoplax adhaerens* | B3S3R6 | Amorphea/Animals | 0.96 |
| *Trichoplax adhaerens* | B3RJ73 | Amorphea/Animals | 0.73 |
| *Monosiga brevicollis* | A9VCG8 | Amorphea/Choanoflagellates | 0.96 |
| *Salpingoeca rosetta* | F2U687 | Amorphea/Choanoflagellates | 0.87 |
| *Amoeboaphelidium occidentale* | KAI3661569.1 | Amorphea/Fungi | 0.50 |
| *Batrachochytrium dendrobatidis* | F4NVJ0 | Amorphea/Fungi | 0.16 |
| *Candida albicans* | A0A1D8PSP4 | Amorphea/Fungi | 0.99 |
| *Coemansia asiatica* | A0A9W8CLX2 | Amorphea/Fungi | 0.00 |
| *Mucor circinelloides* | S2J9K0 | Amorphea/Fungi | 0.98 |
| *Saccharomyces cerevisiae* | P25846 | Amorphea/Fungi | 0.97 |
| *Ustilago maydis* | A0A0D1DQ73 | Amorphea/Fungi | 0.01 |
| *Galdieria sulphuraria* | EME29999.1 | Archaeplastida/Red Algae | 0.12 |
| *Naegleria gruberi* | D2VDU3 | Discoba/Heterolobosea | 0.00 |
| *Andalucia godoyi* | A0A8K0AJE8 | Discoba/Jakobida | 0.98 |
| *Aphanomyces euteiches* | KAH9115059.1 | TSAR/Stramenopiles | 0.57 |
| *Hondaea fermentalgiana* | A0A2R5G5Z1 | TSAR/Stramenopiles | 0.00 |
| *Phytophthora cinnamomi* | KAG6587133.1 | TSAR/Stramenopiles | 0.00 |
| *Saprolegnia parasitica* | XP_12211697.1 | TSAR/Stramenopiles | 0.92 |

Regardless of the origins of the “yeast *MSH1*” gene and its relatives, they are clearly not orthologous to the *CHM/MSH1* gene found in plants. The latter is sometimes referred to as “plant” *MSH1* because it was initially only known to be found in Viridiplantae (Lin et al. 2007; Ogata et al. 2011). More recent database searches have shown that “plant” *MSH1* is also present in various other lineages across the tree of life (Figure 3b), including some protists that have acquired plastids via secondary endosymbiosis, a small clade of Gammaproteobacteria, and multiple nucleocytoviruses (i.e., “giant viruses” or nucleocytoplasmic large DNA viruses) (Wu et al. 2020), all of which share the GIY-YIG endonuclease domain that distinguishes “plant” MSH1 from the rest of MutS family (Abdelnoor et al. 2006; Ogata et al. 2011; Peñafiel-Ayala et al. 2024). This disjunct phylogenetic distribution (Figure 3b) is clear evidence of HGT. Although the gene tree topology is not well enough resolved to confidently identify the HGT donor for an ancient event that must have occurred prior to the MRCA of all Viridiplantae, nucleocytoviruses are arguably the most likely candidate. These viruses can contain genomes more than 1 Mb in size that encode large suites of genes not typically found in viruses, including those involved in DNA repair processes (Aylward et al. 2021). Nucleocytoviruses have already been identified as the likely donor for the *MutS7* gene that is found in the mitochondrial genomes of octocorals (Bilewitch and Degnan 2011; Ogata et al. 2011; Muthye and Lavrov 2021). Therefore, the presence of “plant” *MSH1* genes in extant nucleocytoviruses (Wu et al. 2020) suggests these viruses may be the ultimate source for this distinctive MutS subfamily. Some *MutS* genes in nucleocytoviruses have been found to encode inteins (Green et al. 2018), which themselves are mobile elements. This observation raises the question as to whether inteins have played any role in HGT of these genes from viruses into eukaryotes. However, we are not aware of any cases in which an extant eukaryotic *MutS* gene currently encodes an intein.

First identified based on its effects on plant growth and development (Redei 1973), the functional role of the “plant” *MSH1* gene has now been studied for over half a century, leading to a greatly improved understanding of its underlying cellular and biochemical mechanisms (Table 2). “Plant” *MSH1* encodes an intact MutS Domain I (Figure 1), and disrupting this gene leads to an increase in point mutation rates in both mitochondrial and plastid genomes (Wu et al. 2020), suggesting that it has MMR function. In addition, *msh1* mutants exhibit extensive rearrangements of mitochondrial and plastid genomes mediated by recombination between small repeats, indicating that MSH1 suppresses such illegitimate or ectopic recombination (Martinez-Zapater et al. 1992; Sakamoto et al. 1996; Abdelnoor et al. 2003; Davila et al. 2011; Odahara et al. 2017; Zou et al. 2022). The presence of a C-terminal GIY-YIG endonuclease domain that distinguishes this protein from other MutS subfamilies (Figure 1) prompted the hypothesis that MSH1 uses an atypical DNA repair mechanism. Specifically, the enzyme has been proposed to induce double-stranded breaks (DSBs) in response to recognized mismatches followed by accurate DSB repair via homologous recombination (HR) (Christensen 2014; Ayala-García et al. 2018; Broz et al. 2022). Recent efforts to recombinantly express and purify MSH1 and its domains are creating the opportunity to test these hypothesized enzymatic activities more directly (Fukui et al. 2018; Peñafiel-Ayala et al. 2024). This work has confirmed the predicted nucleotide binding and enzymatic activities towards recombination intermediates, but it surprisingly did not find preferential binding to mismatched DNA templates by MSH1 (Peñafiel-Ayala et al. 2024). However, enzyme recognition and substrate cleavage can be sensitive to reaction conditions. Recent unpublished studies indicate that MSH1 cleaves mismatches and small indels under a narrow range of reaction conditions (Peñafiel-Ayala et al. Unpublished data).

**Table 2.**
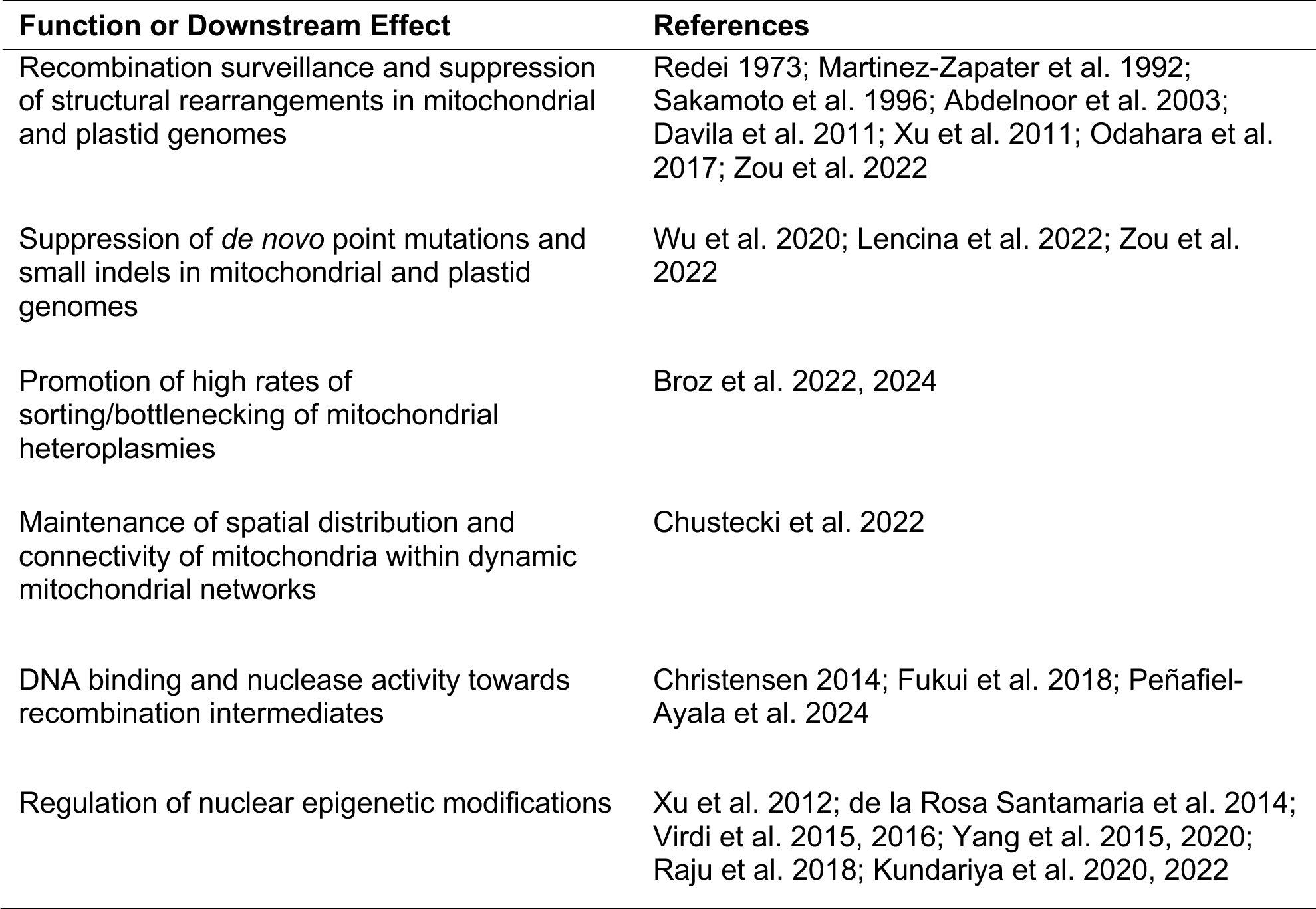
Characterized roles of “plant” MSH1.

In addition to the genomic consequences of disrupting MSH1 function, there are also cellular-level effects on organelle morphology (Sakamoto et al. 1996) and mitochondrial dynamics (Chustecki et al. 2022), including more contacts and a higher degree of connectivity in the mitochondrial networks of *msh1* mutants. Loss of MSH1 activity also appears to slow the “sorting” process by which heteroplasmic point mutations in mitochondria are either lost or spread to fixation (homoplasmy) within an individual (Broz et al. 2022, 2024). An important remaining question is whether such changes are directly related to MSH1 activity or simply reflect indirect consequences of the increases in organelle genome instability in *msh1* mutants. One possibility for a direct relationship between MSH1 activity and the rate of heteroplasmic sorting is that the hypothesized model of DSBs and recombinational repair would be expected to cause gene conversion between different copies of the mitochondrial and plastid genomes, leading to a homogenizing effect that could eliminate heteroplasmies (Broz et al. 2022).

One intriguing development in the study of MSH1 is the identification of diverse phenotypic effects in *msh1* mutants on growth and stress response. Surprisingly, some of these phenotypes do not appear to be mediated by mutations or rearrangements in mitochondrial and plastid genomes but instead may result from changes in nuclear epigenetic markers that show heritability across generations (see references in Table 2). The signaling pathways that govern these epigenetic changes remain to be fully elucidated, as MSH1 is not known to localize to the nucleus. Instead, it has been hypothesized that plastid-mediated signaling in response to disruption of MSH1 activity triggers altered nuclear methylation profiles (Beltrán et al. 2018).

Overall, the gain of MSH1 in Viridiplantae via HGT has had major effects on the genetics of plant organelles. Interestingly, it appears to be central in shaping some of the characteristics that distinguish mitochondrial and plastid genomes in plants from the organellar genomes in other eukaryotes, such as their low mutation rates and highly recombinational nature.

## Ancient Acquisition of MutS1 from Plastids Followed by Fusions with Nuclease-Like Domains and Secondary Losses

Given that members of Viridiplantae have the distinctive “plant” *MSH1* gene that was described in the preceding section, these species have generally been thought to lack bacterial-like *MutS1* genes such as the *MSH1* gene found in some fungi, animals, and other eukaryotes. Although this appears to be true in seed plants, a phylogenetic analysis of the MutS family discovered that other plant lineages contain a cyanobacterial-like *MutS1* gene that was presumably derived from plastid EGT (Hofstatter and Lahr 2021). Although that study described the gene as “chloroplastic *MSH1*,” we will refer to it as *MutS1* to avoid further complicating the already confusing and inconsistent usage of the name *MSH1* in eukaryotes. This gene is found in multiple lineages of land plants, including liverworts, mosses, lycophytes, and ferns (Figure 4), suggesting that the apparent absence of *MutS1* in seed plants is the result of secondary loss. Although it also appears to have been lost from many or most species of green algae, *MutS1* is present in at least some representatives of both chlorophytic algae (e.g., *Cymbomonas tetramitiformis*) and streptophytic algae (e.g., *Zygnema circumcarinatum*). Multiple species of red algae and the glaucophyte *Cyanophora paradoxa* also contain cyanobacterial-like *MutS1* genes. Although phylogenetic analysis does not provide robust support for these genes forming a single monophyletic clade (Figure 4), the presence of a cyanobacterial-like *MutS1* in representatives of all major lineages of Archaeplastida strongly suggests an ancestral origin via plastid EGT.

**Figure 4.**
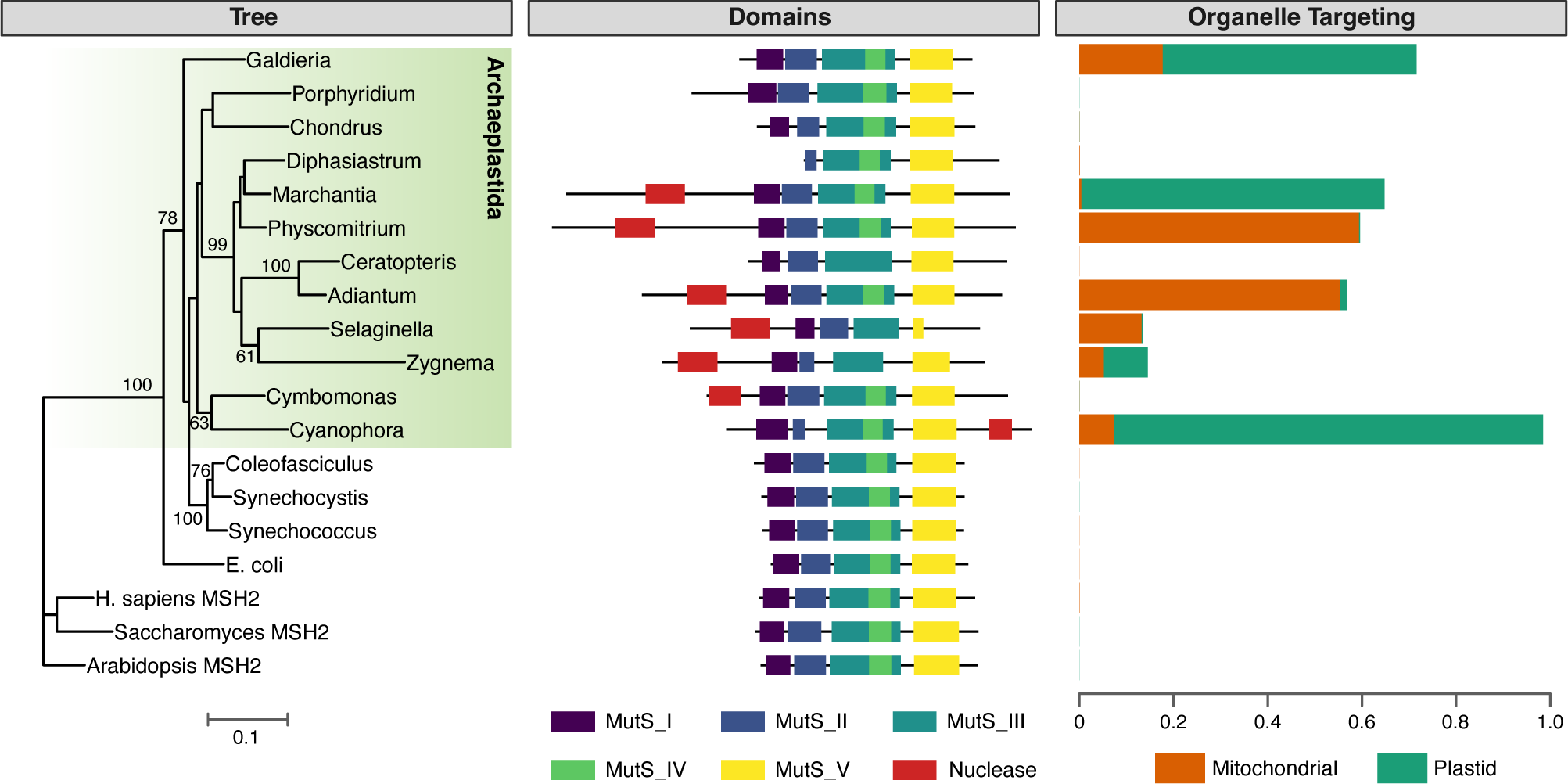
A *MutS1*-like gene in plants. The major Archaeplastida lineages (glaucophytes, red algae, and Viridiplantae) all have representatives with a gene that shows similarities to cyanobacterial *MutS1*, although this gene appears to be absent from seed plants. The maximum-likelihood phylogenetic tree (left) is based on MutS1 protein sequences, using MSH2 for rooting. Bootstrap values are reported for clades with >50% support. The summary of domain architectures (center) reports the classic configuration of MutS1 domains, as well as fusion with an N-terminal DPD1 protein in many representatives of the Viridiplantae lineage or a C-terminal domain with similarities to a Mrr_cat restriction endonuclease domain in *Cyanophora* (but see main text for reasons to be skeptical that this domain functions as a nuclease in *Cyanophora*). Organelle targeting values (right) reflect the probability of subcellular localization to the mitochondria or plastids based on computational predictions by TargetP v2.0 (Armenteros et al. 2019). Many (but not all) MutS1 proteins in the Archaeplastida lineage have some evidence of mitochondrial and/or plastid localization. As expected, none of the bacterial or MSH2 sequences have any predicted organelle targeting. See Supplementary Text S1 for information on tree reconstruction, domain prediction, and protein targeting analysis.

The MutS1 proteins in plants retain the full set of MutS1 domains, including Domain I, which is generally involved in mismatch recognition, and the structural elements in Domain V that are necessary for ATP hydrolysis and conformational changes for mismatch processing (ATP binding site, Walker A motif, Walker B motif, and the signature of a *trans* composite active site). In most species, the retained elements include an anciently conserved FXE motif (Phe36 and Glu38 in *E. coli*; Figure 5b). Mutation of these residues to Ala results in enzymes unable to recognize mismatches (Yamamoto et al. 2000; Schofield et al. 2001). However, in the streptophytic green alga *Zygnema circumcarinatum* and ferns (*Ceratopteris* and *Adiantum*), substitutions have eliminated one or both of these residues (Figure 5a), suggesting that the mismatch recognition domain is compromised in these species.

**Figure 5.**
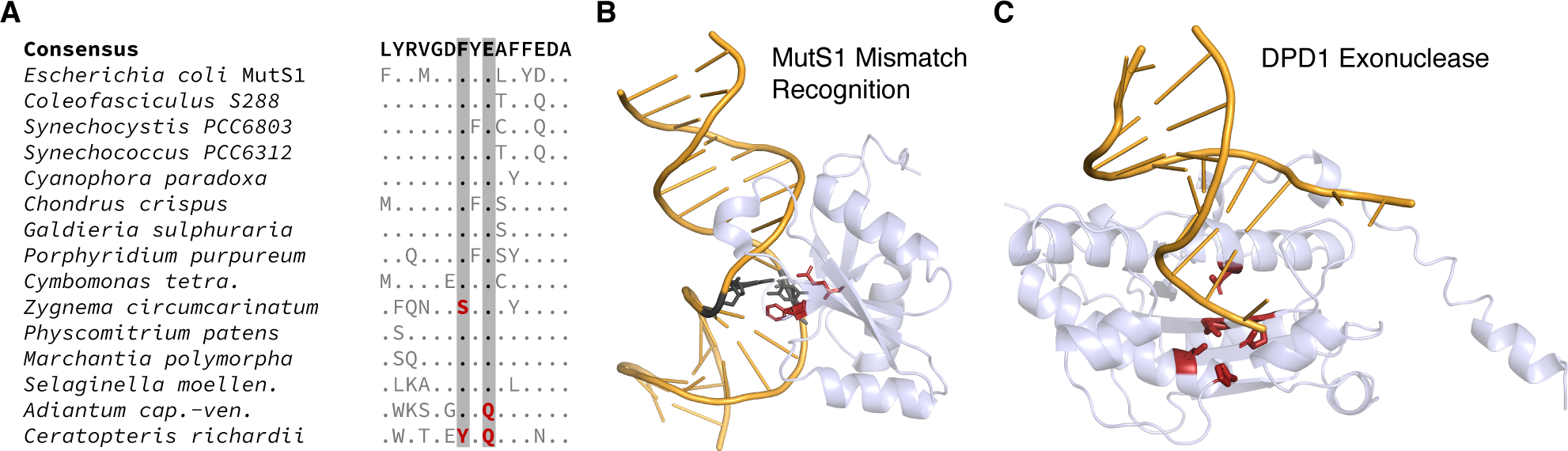
Structural features of cyanobacterial-like MutS1 proteins in plants. (A) Aligned motif within MutS1 Domain I, highlighting the Phe36 and Glu38 residues that are required for mismatch recognition. Only variants relative to the consensus sequence are shown. The substitutions (shown in red) at these positions in the streptophytic alga *Zygnema* and the ferns *Adiantum* and *Ceratopteris* suggest that the MutS1 proteins in these species may have compromised MMR activity. (B) Structure of *E. coli* MutS1 MMR domain (light blue) in complex with dsDNA (orange) with a mismatched G-T base-pair (highlighted in black) from PDB accession 7AI6. The Phe36 and Glu38 residues (highlighted in red) directly interact with this mismatch. The highlighted nucleotides and amino acid residues are also shown in ball-stick representation. (C) Structural model of the N-terminal DPD1 domain (light blue) from the *Physcomitrium patens* MutS1 protein superimposed on the dsDNA substrate (orange) from the crystal structure of TREX2 (PDB 6A47). Predicted active-site residues (Asp276, Glu278, Asp370, His431, and Asp436) are highlighted in red and shown in ball-stick representation. The identity of these residues is conserved in all sampled plant DPD1-MutS1 proteins.

Interestingly, plant MutS1 proteins have evolved accessory domains (Figure 4). Most copies of the gene in Viridiplantae are exceptionally long, in some cases encoding proteins that are approximately twice the size of typical MutS1. In Viridiplantae, many of these proteins have a large N-terminal extension that contains a region orthologous to the angiosperm 3′-5′ exonuclease DPD1 (Defective in Pollen DNA Degradation 1). In *Arabidopsis*, DPD1 (AT5G26940) is expressed in pollen and senescing leaves where it degrades mitochondrial and plastid DNA (Matsushima et al. 2011; Tang and Sakamoto 2011; Takami et al. 2018). In many green algae that appear to have lost the MutS1 protein such as *Micromonas*, *Ostreococcus*, and *Volvox*, a DPD1-like exonuclease is still present and instead fused to the helicase RecG, another DNA binding protein that is targeted to mitochondria and plastids (Odahara et al. 2015; Wallet et al. 2015). The chlorophyte *Cymbomonas tetramitiformis* is distinct in harboring both MutS1 and RecG proteins with DPD1-like fusions. DPD1 is evolutionarily related to proofreading 3′-to-5′ exonucleases like the epsilon subunit of bacterial DNA polymerases (DnaQ) or the editing domain of family-A DNA polymerases (Shevelev and Hübscher 2002; Matsushima et al. 2011; Tang and Sakamoto 2011; Takami et al. 2018). Dali searches (Holm 2022) using the structural models of the DPD1 domain from *Physcomitrium patens* suggest that this protein is highly similar to Three-prime Repair Exonucleases 1 and 2 (TREX1 and TREX2), which belong to the ubiquitous DEDDh exonuclease family and are the main 3′-to-5′ exonucleases in animals (Mazur and Perrino 1999; Cheng et al. 2018). The catalytic residues of TREX proteins are conserved in DPD1 and all sampled DPD1-MutS1 proteins. The TREX active site consists of five amino acids corresponding to Asp276, Glu278, Asp370, His431, and Asp436 in *Physcomitrium patens* DPD1 (Figure 5c). Thus, plant DPD1-MutS1 enzymes harbor a putatively active DPD1 module predicted to display the same biochemical properties as observed in *Arabidopsis* (Matsushima et al. 2011; Tang and Sakamoto 2011; Takami et al. 2018).

In contrast to those found in Viridiplantae, MutS1 proteins in red algae more closely resemble the standalone MutS1 from bacteria without large extensions or predicted nuclease domains. The glaucophyte *Cyanophora paradoxa* also lacks the N-terminal DPD1 fusion, but it has a C-terminal extension (Figure 4) with weak similarity (bitscore: 39.5; E-value 0.0036) to a restriction endonuclease domain (Conserved Domain Database cl46443: Mrr_cat superfamily). Whether this *Cyanophora* domain has any actual nuclease activity is difficult to discern given the lack of closely related protein sequences. Indeed, nuclease activity may be unlikely given the lack of some important catalytic residues. Structure-function studies have shown that Mrr family members harbor three motifs that are highly conserved, with two Asp residues in Motif II and a Lys residue in Motif III being essential for activity (Smith et al. 2009). This signature is not present in the *Cyanophora* MutS1-like sequence, which has an Arg residue instead of Lys in Motif III and exhibits a deletion of one amino acid between the two conserved Asp residues from Motif II. The distance between carboxylates is often crucial for the enzymatic activity, and the catalytic Lys is absolutely conserved in several nucleases (Aravind et al. 2000). Therefore, despite the sequence similarity to the Mrr_cat superfamily, this domain potentially lacks nuclease activity. Regardless of the role of the C-terminal extension of *Cyanophora* MutS1, the variation in domain architecture within this one lineage highlights the remarkable capacity of MutS proteins to acquire accessory domains that presumably expand or alter their functions.

In typical MMR pathways, MutS1 recruits its partner MutL upon mismatch recognition, which coordinates protein-protein interactions and can provide endonuclease function in the repair process (Guarné 2012). Of the major Archaeplastida lineages that have a *MutS1* gene, only red algae also have a cyanobacterial-like *mutL* gene (Hofstatter and Lahr 2021). Assuming that this *mutL* gene was originally gained via plastid endosymbiosis and EGT, one intriguing possibility is that its loss in Viridiplantae and glaucophytes is related to the gain of MutS1 accessory domains with putative nuclease activity or other functions in these lineages (Figure 4).

To our knowledge, no study has yet investigated the function of the cyanobacterial-like MutS1 proteins in plants. There are multiple reasons to infer that they play a role in plastids and/or mitochondria, including their apparent plastid origins via EGT, *in silico* predictions of N-terminal transit peptides that mediate organelle targeting (Figure 4), and the fusion to DPD1, which is known to function in mitochondria and plastids in angiosperms (Matsushima et al. 2011; Takami et al. 2018). Although some Viridiplantae MutS1 proteins lack any predicted transit peptide (e.g., *Diphasiatrum* and *Ceratopteris*; Figure 4), it is not clear whether those are true truncations or just incomplete gene models in genome annotations, as they also lack the N-terminal DPD1 fusion. Given that the typical MutS1 domain architecture remains intact in most of these MutS1 proteins, it is tempting to speculate that they play a role in plastid and/or mitochondrial MMR, as least in cases where the FXE mismatch recognition motif is present (Figure 5a). However, it is surprising that these proteins are found in many Viridiplantae lineages that also contain the organelle-targeted “plant” MSH1, which similarly has domains related to MMR (Figure 1a) (Abdelnoor et al. 2003; Peñafiel-Ayala et al. 2024). How the functions of these two proteins might interrelate in land plant taxa such as liverworts, mosses, lycophytes, and ferns is a key question. The association with the DPD1 exonuclease is also intriguing. The identified role of DPD1 in degrading organellar DNA in angiosperms (Matsushima et al. 2011; Tang and Sakamoto 2011; Takami et al. 2018) implies its association with other proteins. For instance, this enzyme is not expected to degrade DNA molecules without linear ends and may need an endonuclease for efficient processing. From the perspective of MutS1 function, the DNA strand breaks introduced during MMR are typically “chewed back” by exonuclease activity, so one possibility is that the fused DPD1 provides this activity. Another possibility is that DPD1-MutS1 regulates recombination with DPD1 acting on the exposed ends of invading strands in recombination intermediates. Genetic and biochemical studies will be needed to functionally characterize the cyanobacterial-like *MutS1* genes in plants and test these speculative hypotheses. The moss *Physcomitrium* may be the most tractable genetic system for investigations of native function, given the absence of these genes in model systems such as *Chlamydomonas* and *Arabidopsis* (along with all other angiosperms).

## Ancient Acquisition of MutS2 from Plastids Followed by Duplication Prior to the Diversification of Viridiplantae

Within eukaryotes, bacterial-like MutS2 proteins are found exclusively in species that contain plastids, and phylogenetic analysis shows grouping of cyanobacterial MutS2 and nuclear-encoded MutS2 within Archaeplastida, suggestive of ancient EGT from the plastid (Figure 2b) (Lin et al. 2007; Ogata et al. 2011). Subsequent duplication of the *MutS2* gene (Lin et al. 2007) resulted in copies referred to as *MutS2A* (*Arabidopsis* AT5G54090) and *MutS2B* (*Arabidopsis* AT1G65070). Both copies are widely conserved across Viridiplantae, indicating that this duplication preceded the diversification of this lineage (Figure 2b). The ancient nature of this duplication is reflected in their sequence divergence (less than 40% amino acid identity between the two *Arabidopsis* copies). Only a single copy of this gene appears to be present in glaucophytes and red algae. However, the deep splits within the *MutS2* gene tree are not well-resolved, so it cannot be ruled out that the gene duplication event occurred earlier in the evolution of the Archaeplastida and was followed by losses of one copy in the lineages outside Viridiplantae.

To our knowledge, the functions of plant MutS2 proteins have not been previously investigated, and work has only recently been initiated to characterize their role. Nevertheless, some inferences and hypotheses can already be made based on their subcellular targeting in plants and known functions of their bacterial homologs. Both MutS2A and MutS2B contain transit peptides that target them to plastids (Abdelnoor 2004; Carrie et al. 2009; Olinares et al. 2010; Huang et al. 2013), where they have been specifically detected in the nucleoids (Majeran et al. 2012). MutS2B shares considerable structural similarity to bacterial MutS2, which includes a C-terminal extension that folds into three structural modules: a large alpha-helix coiled coil, a KOW domain, and a domain known as Smr (small MutS-related) that has been associated with endonuclease activity (Cerullo et al. 2022). In contrast, MutS2A lacks the C-terminal Smr domain (Figure 1a). In bacteria, MutS2 proteins form homodimers (Figure 1b), but the presence of ancient duplicates in Viridiplantae raises the possibility that MutS2A and MutS2B heterodimerize. Thus, determining the protein-protein interactions and dimerization state of plant MutS2 proteins is an important goal in characterizing their function.

Because MutS2 proteins lack Domain I (Figure 1a), which is responsible for mismatch recognition, they are thought to instead function in HR. Supporting this idea, bacterial MutS2 binds recombination intermediates such as Holliday junctions and D-loops using its core/clamp (Domains III and IV) (Kang et al. 2005; Pinto et al. 2005; Fukui et al. 2022). Interestingly, however, bacterial MutS2 has been shown to have contrasting roles in HR, either suppressing it in *Thermus thermophilus* (Fukui et al. 2008) and *Heliobacter pylori* (Pinto et al. 2005; Damke et al. 2015) or promoting it in *Bacillus subtilis* (Burby and Simmons 2017). Examples of anti-recombination activity have been attributed to the Smr endonuclease domain, which can cleave and resolve recombination intermediates (Fukui et al. 2004, 2007, 2008; Jeong et al. 2012; Zhang et al. 2014; Damke et al. 2015). In contrast, the promotion or stabilization of recombination intermediates is more reminiscent of the activity found in eukaryotic MSH4/MSH5 (Ross-Macdonald and Roeder 1994; Hollingsworth et al. 1995; Snowden et al. 2004), both of which have a domain architecture that is similar to MutS2 but lacks the Smr domain (Figure 1a). As Viridiplantae contain MutS2 homologs both with and without the Smr domain, it is unclear whether these proteins might act to promote or suppress recombinational activity (or neither) in plastids. The role of MutS2B may also depend on key sequence variants in the Smr domain that have been implicated in functional differences among Smr-containing proteins (see below).

Recent work in *Bacillus subtilis* uncovered an additional function of MutS2 in quality control of translation through ribosomal rescue (Cerullo et al. 2022; Park et al. 2024). MutS2 is recruited to collided ribosomes via its C-terminal Smr and KOW domains (Park et al. 2024) and binds the leading ribosome using its clamp domain (IV) (Cerullo et al. 2022). An ATP-dependent conformational change then drives splitting of the leading ribosome, releasing it from the mRNA and initiating the process of ribosomal quality control (Cerullo et al. 2022). It was initially presumed that the Smr domain of MutS2 would cleave the polysome-associated mRNA (similar to the function of *E. coli* SmrB), but no evidence was found for this mechanism in subsequent investigations (Park et al. 2024).

Indeed, the nuclease activity of Smr domains is subject to some uncertainty and controversy, and there appears to be functional variation among Smr-containing proteins. In *Thermus thermophilus*, the residues Asp669/Arg671 and His701 were found to form a nuclease active site and be essential for catalysis as part of DXH/R and HGXG motifs, respectively (Fukui et al. 2008; Fukui and Kuramitsu 2011). However, mutagenesis studies implicate the identity of the residue between the two conserved Gly residues in the HGXG domain as critical for catalysis. This residue is a Thr in PPR-SMR proteins, an Arg in Cue2, and a Lys in bacterial MutS2, all amino acids that can contribute to catalysis (Zhou et al. 2017; D’Orazio et al. 2019). In plant MutS2B, this residue is a Met, which lacks potential to be involved in catalysis. Furthermore, structure-function studies of the *E. coli* SmrB protein concluded that the DXH motif is solely responsible for nuclease activity (Saito et al. 2022). Accordingly, the lack of Smr nuclease activity in *Bacillus* MutS2 was attributed to its DXR variant of the DXH motif being unable to promote mRNA cleavage (Park et al. 2024). Thus, while proteins like Cue2 that carry nuclease-active Smr domains may rescue stalled ribosomes by cleaving mRNA, proteins harboring nuclease-inactive Smr domains like some bacterial MutS2 enzymes and potentially plant MutS2B may function by “ribosome-splitting” (D’Orazio et al. 2019; Park et al. 2024). More generally, some Smr domains that lack nuclease activity (including distant relatives such as ALBA proteins) may act as RNA chaperones (Glover et al. 2020; Tong et al. 2022).

Other work has implicated MutS2 in cellular responses to oxidative stress (Wang et al. 2005; Zhang et al. 2014; Wang and Maier 2017). In both *Helicobacter pylori* and *Deinococcus radiodurans*, *mutS2* mutants are more sensitive to oxidative stress than wild type cells and preferentially bind damaged DNA containing 7,8-dihydro-8-oxoguanine (8-oxoG) (Wang et al. 2005; Zhang et al. 2014). In *Helicobacter pylori*, DNA binding is associated with a motif directly upstream of the Smr domain that appears to be unique to that species. Overall, the remarkable diversity of bacterial MutS2 functions raises intriguing possibilities that MutS2 plays one or more similar roles in plants.

## Summary

The expanded set of MutS proteins in plants relative to other eukaryotic lineages is the result of a combination of gene duplication, HGT, and EGT (Figure 6). The propensity of these proteins to gain and lose accessory domains, including many with putative nuclease activity, has apparently contributed to their functional diversification. However, there is much to learn about the actual role of these proteins in plants, especially MutS1 and MutS2, which remain almost entirely uncharacterized. Notably, all of the proteins that originated in plants due to HGT or EGT (MSH1, MutS1, and MutS2) have known or hypothesized roles in mitochondria and/or plastids, indicating outsized contributions to shaping the distinctive features of plant organelle genetics, such as low mutation rates, extensive recombinational activity, and massive levels of protein expression.

**Figure 6.**
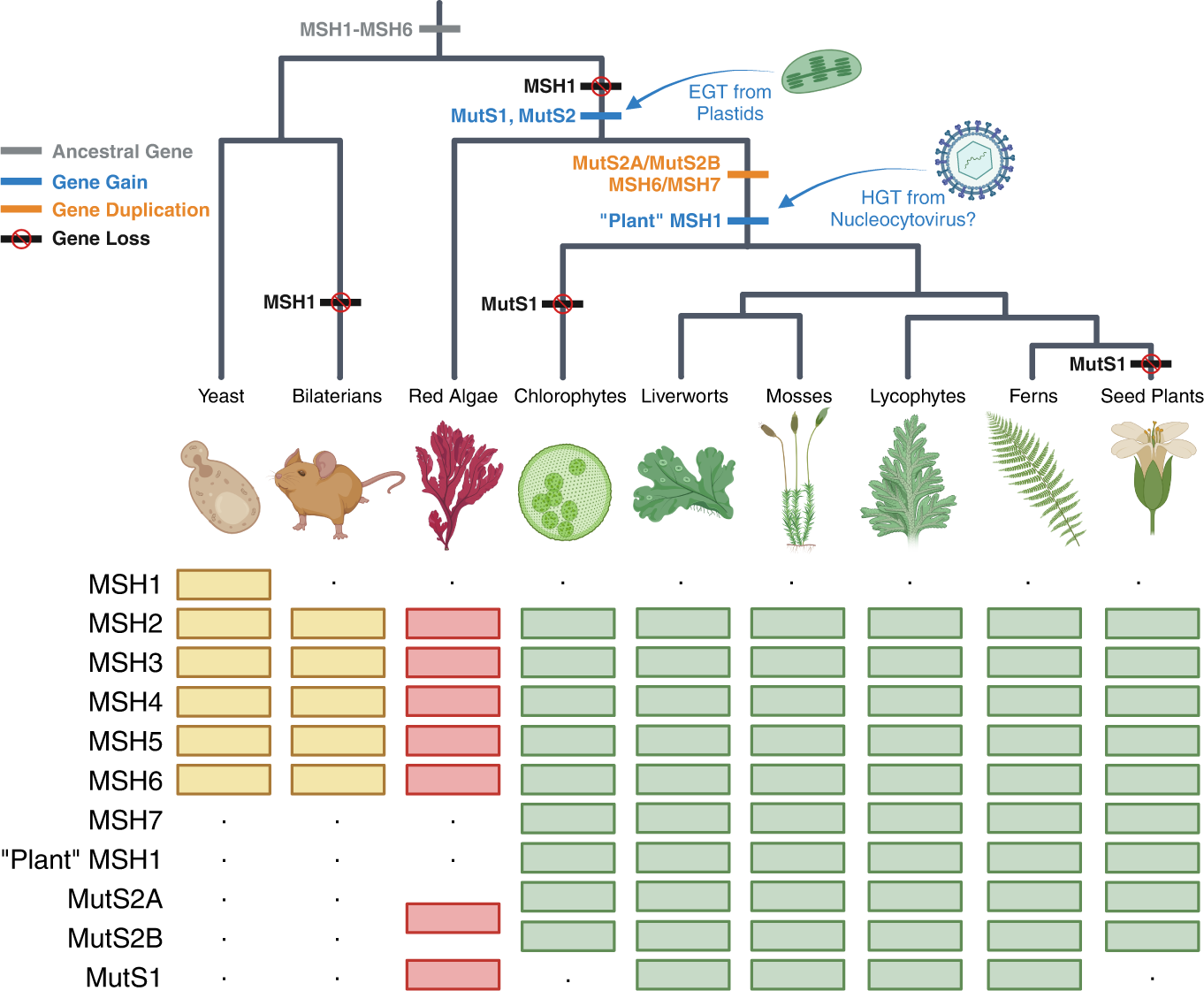
History of *MutS* gene family expansion in plants with the inferred timing of gene gains, losses, and duplications indicated on the tree. Gene presence or absence in a lineage is indicated by boxes and dots, respectively. In general, phylogenetic reconstructions of deep splits in MutS gene trees are not well supported or are prone to long-branch artefacts, so alternative scenarios are possible. For example, the MSH6 and MutS2 duplications could have occurred earlier and been followed by losses in lineages outside of Viridiplantae. A MutS1 loss is indicated for chlorophytes because it appears to be absent from most sampled species in this lineage. However, the presence of MutS1 in *Cymbomonas tetramitiformis* indicates that it is not entirely absent from all chlorophytes. This figure was generated with Biorender.

## Acknowledgements

This work was supported by the National Institutes of Health (R35GM148134, T32GM132057), the National Science Foundation Graduate Research Fellowship Program, and CONAHCYT-Fronteras de la Ciencia (170713).

## Supplementary Text S1

*Maximum Likelihood Phylogenetic Inference*. Following exploratory taxon-restricted BLASTP searches of the NCBI nr database and Orthofinder (Emms and Kelly 2019) clustering of sequences for target taxa, amino acid sequences for identified MutS proteins were aligned with the L-INS-i algorithm in MAFFT v7.520 (Katoh and Standley 2013) and trimmed with trimAl v1.4.rev22 build[2015-05-21] (Capella-Gutierrez et al. 2009), retaining alignment positions with a minimum of 80% non-gapped sequences (-gt 0.8). Selection of optimal models of sequence evolution and maximum likelihood tree searches were performed using IQ-TREE v2.3.0 (Mar 14 2024 build) with 1000 parametric bootstrap replicates (Minh et al. 2020). Amino acid sequence alignments (both trimmed and untrimmed) and Newick-formatted phylogenetic trees are available via GitHub (https://github.com/dbsloan/plant_MutS_expansion).

*Mitochondrial and Plastid Targeting Predictions*. Prediction of subcellular localization to mitochondria and/or plastids was performed with TargetP v2.0 (Emanuelsson et al. 2000). The “Plant” option was selected for plant-specific datasets.

*Protein Domain Predictions*. InterProScan (Blum et al. 2021) was used to search for Pfam domains in each protein sequence, including MutS Domains I-V (PF01624, PF05188, PF05192, PF05190, and PF00488) and associated nuclease domains (PF00929, PF01541, PF01713). In some cases, Pfam MutS Domain III was not detected, but a hit to SMART Domain SM00533 was detected, which was then used to define the boundaries of MutS Domain III. In addition, sequences were searched against the NCBI Conserved Domain Database with the CD-SEARCH web tool (Marchler-Bauer and Bryant 2004), which identified an Mrr_cat domain hit in the C-terminal extension of *Cyanophora* MutS1.

*Protein Structural Modeling and Alignment*. Relevant protein structures were obtained from the Protein Data Bank as indicated in the corresponding figure legends and visualized with PyMol v3.0.3. A BLASTP search using the DPD1 amino acid sequence (residues 259 to 494) of *Physcomitrium patens* MutS1 identified TREX2 as the experimentally solved protein structure that shares the largest percentage of amino acid identity. The DPD1 domain of *Physcomitrium patens* MutS1 was used to generate an AlphaFold2 structural model specifying as a template the crystal structure of TREX2 (Perrino et al. 2005; Cheng et al. 2018). The 3′-overhang DNA from the TREX2-DNA crystal structure was superimposed onto the resulting DPD1 structural model.

